# Evaluating the role of inhibiting the biosynthesis of estrogens in the sex-specific antidepressant-like effects of ketamine in rats

**DOI:** 10.1101/2023.03.29.534716

**Authors:** Sandra Ledesma-Corvi, Jordi Jornet-Plaza, M. Julia García-Fuster

## Abstract

Ketamine has been recently approved to treat resistant depression; however preclinical studies showed sex differences in its efficacy. Sex steroids, such as estrogen and testosterone, both in the periphery and locally in the brain, are regarded as important modulators of these sex differences. Therefore, the present study evaluated the role of inhibiting estrogen biosynthesis with letrozole, an aromatase inhibitor that catalyzes the conversion of androgen into estrogen, in the differential antidepressant-like-response induced by ketamine with sex. We performed several consecutive studies in adult Sprague-Dawley rats to evaluate potential sex differences in the antidepressant-like effects of ketamine (5 mg/kg, 7 days, i.p.), letrozole (1 mg/kg, 8 days, i.p.) and their combination (letrozole pre-treatment 3 h before ketamine). Acute and repeated antidepressant-like responses were ascertained in a series of behavioral tests (forced-swim, novelty-suppressed feeding, two-bottle choice for sucrose preference). The main results proved clear sex differences in the antidepressant-like response induced by ketamine, which was observed following a repeated paradigm in adult male rats, but rendered inefficacious in female rats. Moreover, decreasing estrogen production with letrozole induced on itself an antidepressant-like response in female rats, while also improved ketamine’s response in male rats (i.e., quicker response, only after a single dose). Interestingly, both the antidepressant-like effects induced by ketamine in male rats or letrozole in female rats persisted over time up to 65 days post-treatment, suggesting long-term sex-directed benefits for these drugs. The present results demonstrated a sex-specific role for inhibiting estrogen biosynthesis in the antidepressant-like response induced by ketamine in male rats. Moreover, letrozole presented itself as a potential antidepressant for females with persistent effects over time. Clearly, estrogen production is key in modulating, in a sex-specific manner, affective-like responses and thus deserve further studies.

## Introduction

Since esketamine, the S-enantiomer of ketamine, was approved in 2019 by the FDA (https://www.accessdata.fda.gov/drugsatfda_docs/label/2019/211243lbl.pdf) for treatment-resistant depression it has been a hot topic in several international neuropsychopharmacology meetings (e.g., 35th ECNP Congress, 2022). Its great clinical outcome for depressed patients that are resistant to at least two other pharmacological options, confront with the scarce knowledge regarding the potential long-term impact of its repeated treatment (e.g., safety and tolerability; recently reviewed by [1]). In this context, a great effort has been placed in the last years attempting to further characterize ketamine’s antidepressant-like actions, beyond its initial binding as a non-competitive antagonist to the N-Methyl-D-Aspartate (NMDA) receptor (e.g., [1–3]).

Additionally, further defining potential sex differences in ketamine’s therapeutic actions is a priority since targeted treatments might be needed for each sex; in fact, both prior clinical and preclinical reports suggest different responses for males and females, with even contradictory data regarding which sex might benefit the most from the treatment. For example, when analyzing recent reviews with clinical data, no consensus could be reached, since ketamine is sometimes presented as more effective for males (e.g., [4]), but others for females (e.g., [1]), or even with no significant sex differences in its therapeutic effects (e.g., [5]). Similarly, results from preclinical studies are also inconsistent, with improved responses for ketamine for either male or female rodents, that seemed to depend on the dose administered [6–8], the interaction with prior early-life stress exposure [9], and/or the time-duration of the response (e.g., [10]). These sex differences in efficacy reinforce the need of including sex as a biological variable in all preclinical studies [11–13] and might be explained by disparities in the pharmacokinetics and pharmacodynamics effects of the drug [14,15], which could in turn be influenced by a variety of factors including the level of sex steroids, such as estrogen and testosterone, both in the periphery and locally in the brain (e.g., [1]).

In this context, recent studies not only supported a possible involvement of estrogen in the sex-differences reported in the pathogenesis of depression (e.g., [16]), but also in the sex-specific changes in efficacy of certain antidepressants (e.g., [16–17]). However, how the levels of estrogen might be affecting the antidepressant-like response of ketamine is still unknown. One pharmacological tool to evaluate the potential role of estrogen biosynthesis in ketamine’s actions is letrozole, an aromatase inhibitor that prevents estrogen biosynthesis via androgen aromatization (e.g., [18]), which was previously used to evaluate if estrogens would interfere in the antidepressant-like response of fluoxetine in rats [19]. Against this background, the present study utilized male and female Sprague-Dawley rats to characterize the role of inhibiting estrogen biosynthesis with letrozole in the antidepressant-like response induced by ketamine.

## Materials and Methods

### Animals

A total of 145 adult Sprague-Dawley rats (72 male and 73 female) were used in 3 consecutive studies (see Fig. 1). All rats were bred in the animal facility at the University of the Balearic Islands and were housed in standard cages (2-4 rats per cage) with *ad libitum* access to a standard diet and tap water in a controlled temperature (22 °C) and humidity (70%) vivarium (12:12 h light/dark cycle). Procedures were performed under the ARRIVE guidelines [20] and following the EU Directive 2010/63/EU of the European Parliament and of the Council, after approval by the Local Bioethical Committee (University of the Balearic Islands) and the regional Government (Conselleria Medi Ambient, Agricultura i Pesca, Direcció General Agricultura i Ramaderia, Govern de les Illes Balears). Rats were used to being handled by the experimenter prior to any procedures and body weight was monitored across time. All efforts were made to reduce the number of rats used and their suffering. The specific stages of the estrous cycle were not monitored during the experimental procedures since the cyclicity of females was not part of our research question (see [12]), but also because female rats are not more variable than male rats in neuroscience research due to hormonal periodicity (e.g., [21–22]).

**Fig. 1.**
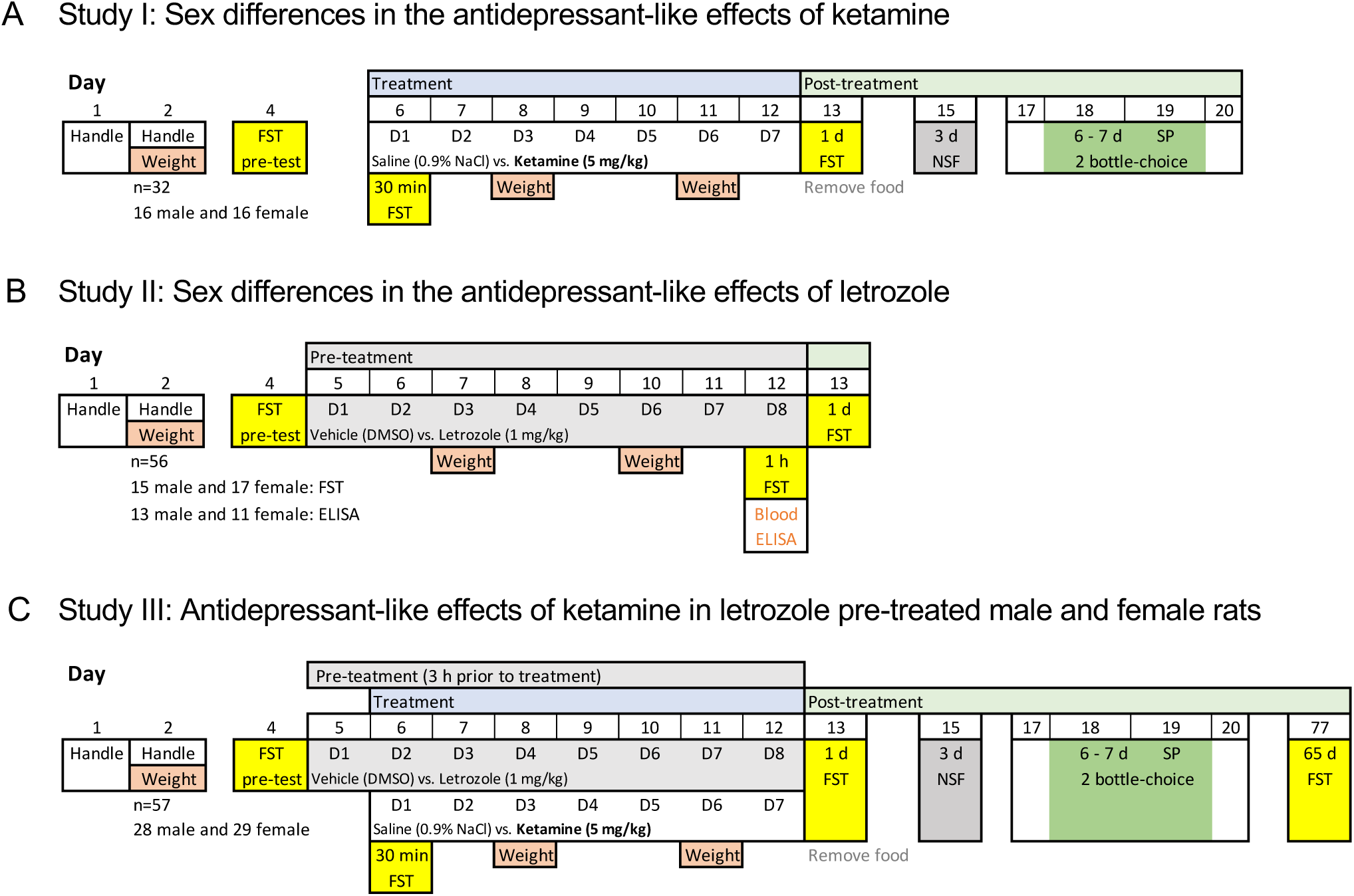
Experimental timeline. **A** Study I: Sex differences in the antidepressant-like effects of ketamine. **B** Study II: Sex differences in the antidepressant-like effects of letrozole. **C** Study III: Antidepressant-like effects of ketamine in letrozole pre-treated male and female rats. FST: forced-swim test; NSF: novelty suppressed feeding; SP: sucrose preference; D: day of treatment, d: day of post-treatment.

### Pharmacological drug treatments

The potential sex differences in the antidepressant-like effects of ketamine were assessed in adult male (n = 16) and female (n = 17) rats that were treated for 7 consecutive days (1 injection per day, i.p., 1 ml/kg) with saline (0.9% NaCl) or ketamine (Richter Pharma, Austria; dose of 5 mg/kg dissolved in saline, selected from [9]) (see Study I; Fig. 1A). To characterize the effects of letrozole (Study II; Fig. 1B), adult male (n = 28) and female (n = 28) rats were treated for 8 consecutive days (1 injection per day, i.p., 1 ml/kg) with vehicle (DMSO) or letrozole (Novartis Pharma, Switzerland, dose of 1 mg/kg, selected from [19]). Some of these rats were exposed to behavioral phenotyping through the forced-swim test (15 male and 17 female) while the rest were used to assess testosterone levels (13 male and 11 female) following the procedures described later on. Finally, we combined both designs to evaluate the role of inhibiting the synthesis of estrogen with letrozole in the differential sex-response induced by ketamine (Study III; Fig. 1C). To do so, male (n = 28) and female (n = 29) rats were pretreated (i.p.) with letrozole (1 mg/kg/day) or vehicle (1 ml/kg/day of DMSO) for 8 consecutive days, followed 3 h later by a daily injection with ketamine (5 mg/kg/day) or saline (1 ml/kg/day of 0.9% NaCl) for 7 consecutive days (days 2-8 of letrozole treatment; see Fig. 1C).

### Hormonal assay

Blood samples were collected during Study II from 13 male and 11 female rats treated with either vehicle or letrozole for 8 consecutive days (blood collection 1 h post injection; see Fig. 1B). To recover plasma, blood samples were centrifuged (4 °C, 15 min at 1500 x g) and then stored at -80 °C. The levels of accumulated testosterone through inhibiting the synthesis of estrogen with letrozole was used as an indicative of hormonal status, and was quantified by a standard ELISA kit (LDN, AR E-8000R, Nordhorn, Germany), according to the manufacturer’s instructions. The sensitivity of the assay was 0.066 ng/ml.

### Forced-swim test

To evaluate the potential antidepressant-like effects of ketamine, letrozole and/or their combination, we first screened the animals under the stress conditions of the forced-swim test [23]. To do so, we followed prior well-established procedures in our group (e.g., 24-26]), in which rats were placed in individual tanks (41 cm high x 32 cm diameter, 25 cm depth) filled with water (25 ± 1 °C) during 15 min (pre-test sessions) followed by 5-min test sessions that were videotaped (see Fig. 1). Test sessions were repeated across time in an attempt to evaluate the progression of the response: acute effects (e.g., 30 min post-injection), repeated effects (e.g., 1 h or 1-day post-treatment) and/or long-term effects (e.g., 65 days post-treatment). Videos were blindly analyzed to determine individual immobility vs. active behaviors (climbing or swimming) for each rat (Behavioral Tracker software, CA, USA).

### Novelty-suppressed feeding test

This test was first described to assess differences in sensitivity to novelty in an anxiogenic-like environment [27], and could be used to score chronic antidepressant-like responses (e.g., [9]). To do so, rats from Studies I and III were food-deprived prior to testing for 48 h, since motivation for food is required (see Fig. 1A and 1C). During testing, which was done 3 days post-treatment, rats were individually placed at one of the corners facing the wall of the arena and were left undisturbed for 5 min in a square open-field arena (60 cm × 60 cm, and 40 cm in high), under housing illumination conditions with three food pellets in the center (e.g., [9, 25]). Sessions were videotaped and the parameters of video analysis that were evaluated for each rat were feeding time (s), and distance traveled (cm).

### Sucrose preference test

This test is used as an indicator of hedonic-like responses (i.e., anhedonia, a core symptom of depression; see [28]), since animals, when given the chance, prefer a sweet solution (e.g., 1% sucrose) over water. Prior to testing, rats from Studies I and III were trained during 24 h to drink from two bottles filled with water that were placed on each side of the housing cage (5 days post-treatment). Then, for 2 consecutive days, rats could either drink from a bottle containing 1% sucrose or another one with water (6–7 days post-treatment; see Fig. 1A and 1C). To avoid the possible preference to a particular side of the cage, bottles were placed in alternate positions. Finally, rats were again exposed for 24 h to two water bottles to make sure no bias for either bottle was present. All bottles were weighted every day to calculate sucrose intake (g/kg) and preference (%) for each cage (groups of 2-4 rats/cage) on days 6 and 7 post-treatment.

### Data analysis and statistics

Data was analyzed with GraphPad Prism, Version 9.4.0 (GraphPad Software, San Diego, CA). In line with the guidelines for displaying data and statistical methods in experimental pharmacology (e.g., [29, 30]), results are presented as mean values ± standard errors of the mean (SEM), and individual symbols are shown for each rat. Each set of data was evaluated either with two-(independent variables: Sex and Treatment or Pre-treatment) or three-way (independent variables: Sex, Pre-treatment and Treatment) ANOVAs. When sex differences emerged, data was also analyzed for each sex separately by two-way ANOVAs (independent variables: Pre-treatment and Treatment) or Student’s *t* test. When appropriate, multiple comparisons were performed by Sidak’s test. The level of significance was fixed at *p* ≤ 0.05.

## Results

### Study I: Sex differences in the antidepressant-like effects of ketamine

Before phenotyping the behavioral responses induced by ketamine, and when only considering potential sex differences in the assays evaluated (effects of Sex; Supplemental Table S1), some significant basal changes emerged for male vs. female rats in all tests that conditioned and/or justified the posterior analysis for each sex separately. For example, there was an overall sex difference in climbing behavior as measured 30 min post a single injection in the forced-swim test (Fig. 2A; Supplemental Table S1: effects of Sex), with female rats climbing less time (-30 ± 13 s, #*p* = 0.031) as compared to male adult rats. Again, sex differences were observed when evaluating distance traveled (cm) in the novelty-suppressed feeding test, with females covering significant more distance (+394 ± 151 cm, ##*p* = 0.003) than male rats (Fig. 2C). Finally, a significant sex difference was also observed for sucrose intake (see Supplemental Table S1), with female rats consuming more sucrose (+3.8 ± 0.6 g/kg, ###*p* < 0.001) than male adult rats (Fig. 2D). Therefore, depending on whether the particular feature of study was or not affected by sex, statistical analyses were performed by two-way ANOVAs or through Students *t*-tests for each sex separately depending on the number of groups evaluated.

**Fig. 2.**
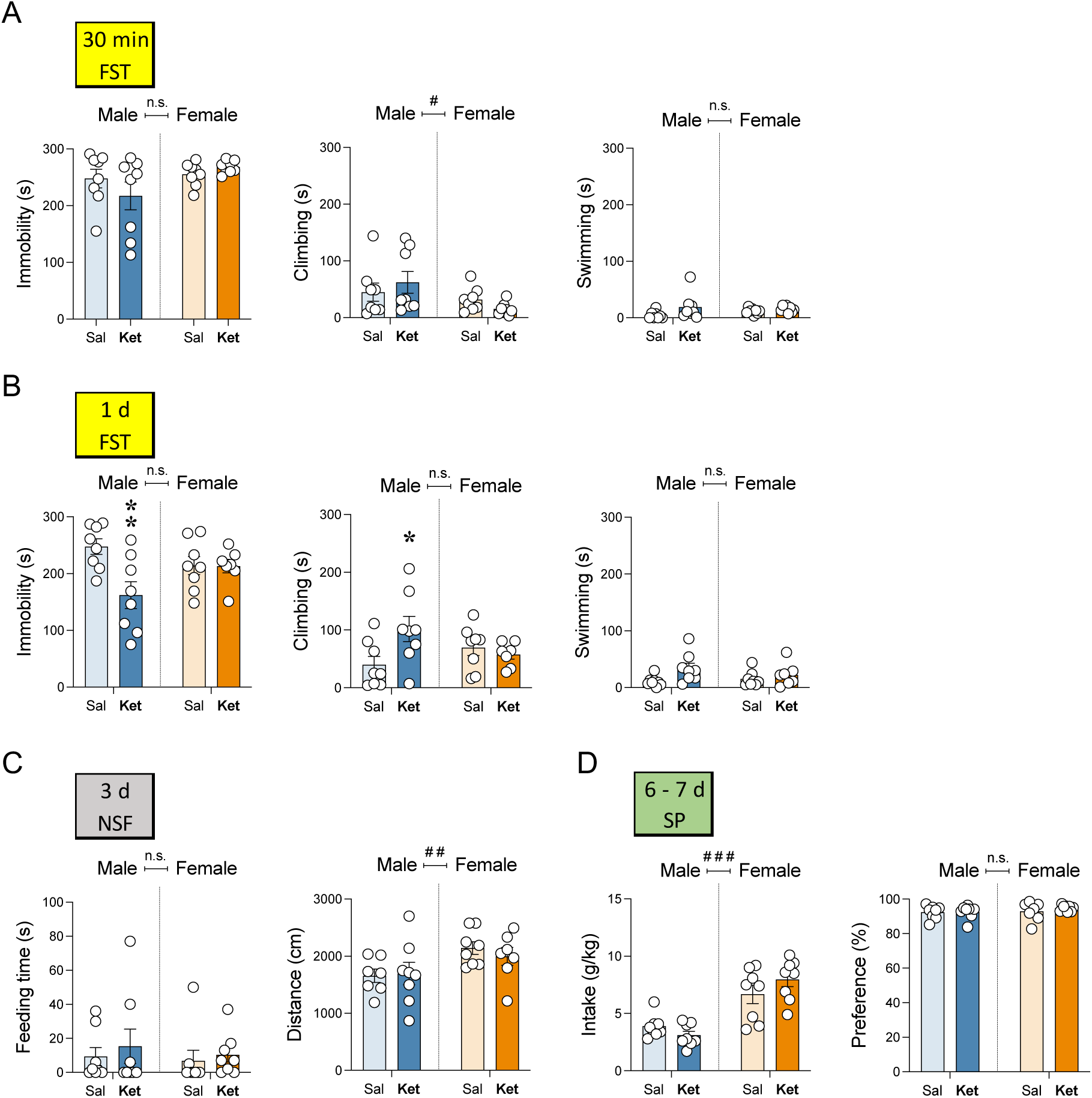
Sex differences in the antidepressant-like effects of ketamine. **A** Acute (30 min post-treatment) and **B** repeated (1-day post-treatment) effects exerted by ketamine exposure in male and female rats in the forced-swim test (FST). Data represent mean ± SEM of the time spent (s) immobile, climbing, or swimming. **C** Repeated effects (3-days post-treatment) exerted by ketamine exposure in male and female rats in the novelty-suppressed feeding test (NSF). Data represent mean ± SEM of the feeding time (s) or distance traveled (cm). **D** Repeated effects (6–7 days post-treatment) exerted by ketamine exposure in male and female rats in the sucrose preference test (SP). Data represent mean ± SEM of sucrose intake (g/kg) or preference (%). **A-D** Individual values are shown for each rat (symbols). Two-way ANOVAs (independent variables: Sex and Treatment) were performed and results are shown in Supplemental Table S1. ^###^*p*<0.001 and ^#^*p*<0.05 when comparing female vs. male rats (effect of Sex). Student’s *t*-tests for each sex separately: \*\**p*<0.01 and \**p*<0.05 vs. same sex saline-treated rats. Sal: saline; Ket: ketamine; Veh: vehicle; LTZ: letrozole.

The acute effect of ketamine was ascertained in the forced-swim test 30 min post a single injection. The results showed that ketamine did not induce changes in immobility, climbing or swimming behaviors in adult male and female rats (Fig. 2A; Supplemental Table S1: lack of significant effects of Treatment or Sex x Treatment interactions). Interestingly, following a repeated 7-day paradigm of ketamine, significant Sex x Treatment interactions emerged, as measured 1-day post-treatment, both for immobility and climbing behaviors (Fig. 2B; Supplemental Table S1). Sidak’s *post-hoc* comparisons revealed that ketamine induced an antidepressant-like effect exclusively in adult male rats, while rendered inefficacious in females (Fig. 2B). Particularly, in male rats, ketamine decreased immobility (-86 ± 23 s, \*\**p* = 0.003) and increased climbing (+62 ± 24 s, \**p* = 0.016) behavior in the forced swim test (Fig. 2B) when compared to saline-treated rats. These effects were not observed later on with other tests also used to characterize the antidepressant-like response. The novelty-suppressed feeding test was performed 3 days post-treatment to measure feeding time (s) and distance traveled (cm) as indicatives of an antidepressant-like response. However, the results showed no significant effects of Treatment, nor significant Sex x Treatment interactions (Fig. 2C; Supplemental Table S1). Similarly, no significant effects of Treatment were observed when measuring 1% sucrose intake (g/kg) or preference (%) in the sucrose preference test in male and female rats 6–7 days post-treatment (Fig. 2D; Supplemental Table S1).

### Study II: Sex differences in the antidepressant-like effects of letrozole

Before phenotyping the behavioral responses induced by letrozole, and as a positive control of the treatment success, we evaluated testosterone levels in plasma (ng/ml), as an indicative of the correct inhibition of estrogen’s biosynthesis by this aromatase inhibitor. As expected, the results showed significant baseline sex differences in testosterone levels (Fig. 3A and Supplemental Table S1), with female rats displaying lower levels (-4.41 ± 1.34 ng/ml of testosterone, ##*p* = 0.004) than males. Interestingly, and given the baseline sex difference in testosterone levels, when the effect of letrozole treatment was evaluated separately for each sex, it only rendered significant for females (+1.89 ± 0.42 ng/ml; *t* = 4.51, df = 9, \*\**p* = 0.002 vs. vehicle-treated female rats), but not for male rats (Fig. 3A).

**Fig. 3.**
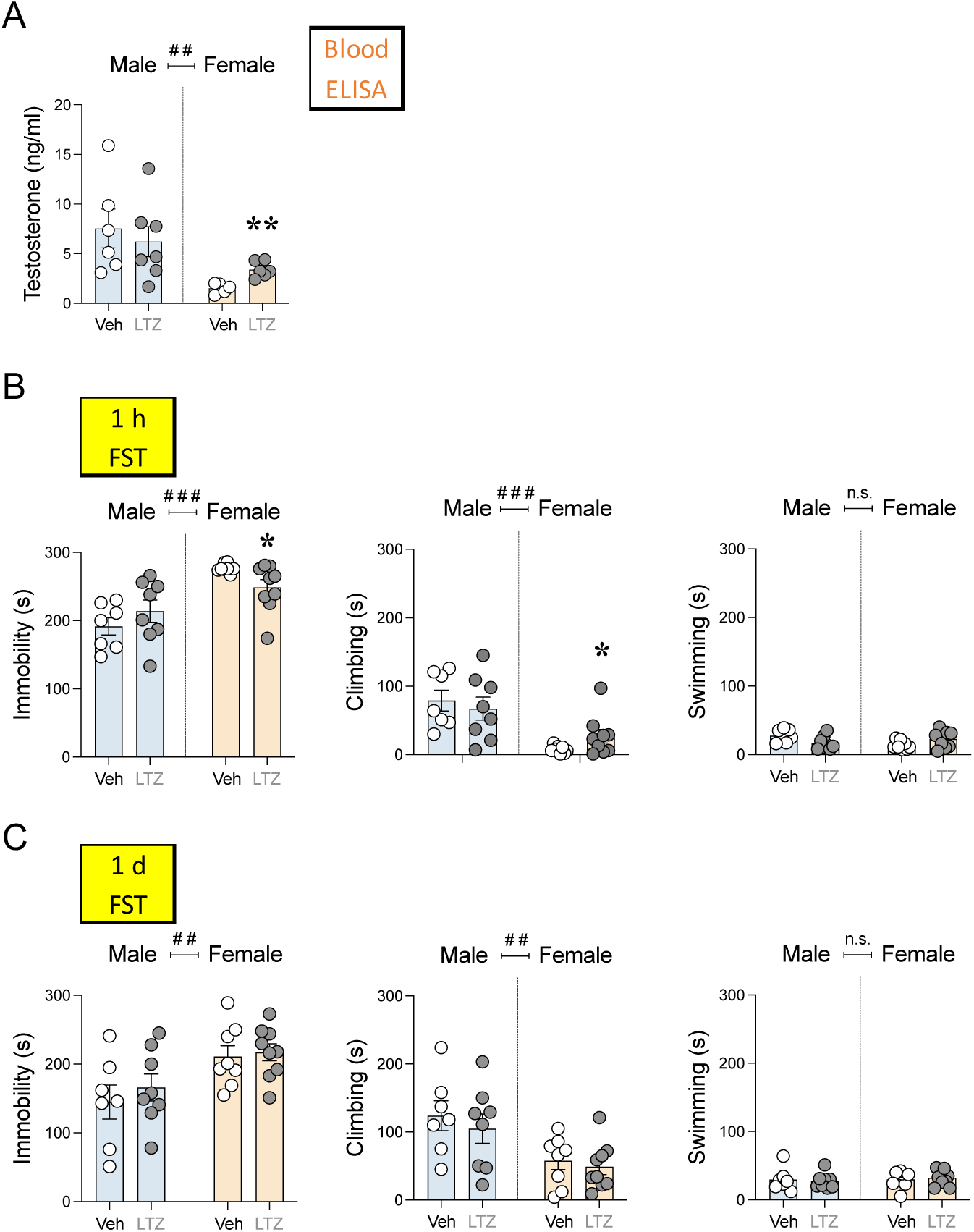
Sex differences in the antidepressant-like effects of letrozole. **A** Repeated effect of letrozole (1 h post-8 days of treatment) on testosterone levels. Data represent mean ± SEM of testosterone levels (ng/ml). **B** Acute (1 h post-8 days of treatment) and **C** repeated (1-day post-8 days of treatment) effects exerted by letrozole exposure in male and female rats in the forced-swim test (FST). Data represent mean ± SEM of the time spent (s) immobile, climbing, or swimming. **A-C** Individual values are shown for each rat (symbols). Two-way ANOVAs (independent variables: Sex and Treatment) were performed and results are shown in Supplemental Table S1. ^###^*p*<0.001 and ^##^*p*<0.01 when comparing female vs. male rats (effect of Sex). \*\**p*<0.01 and \**p*<0.05 vs. same-sex vehicle-treated rats. Veh: vehicle; LTZ: letrozole.

The potential antidepressant-like effect of letrozole was evaluated in the forced-swim test 1 h and 1-day post-repeated treatment (8 days). Significant effects of Sex were observed for almost all data analyzed through two-way ANOVAs (see Supplemental Table S1). Overall, female rats displayed higher immobility (1 h: +60 ± 12 s, ###*p* < 0.001; 1 d: +59 ± 18 s, ##*p* = 0.003) and lower climbing (1 h: -56 ± 12 s, ###*p* < 0.001; 1 d: -61± 17 s, ##*p* = 0.001) than males (Fig. 3B and 3C). Thus, the response of letrozole in the forced-swim test was analyzed for each sex separately through Student’s *t* tests. The results showed that letrozole induced an antidepressant-like effect exclusively in female rats by decreasing immobility (-28 ± 12 s; *t* = 2.27, df = 15, \**p* = 0.038 vs. vehicle-treated female rats), and increasing climbing behavior (+19 ± 11 s; *t* = 1.80, df = 15, \**p* = 0.046 vs. vehicle-treated female rats; one-tailed *p* value) as measured 1 h post-treatment (Fig. 3B). This antidepressant-like effect induced by letrozole in female rats returned to normal 1-day post-treatment (Fig. 3C). No significant effects were observed 1 h or 1-day post-treatment for male rats (Fig. 3B and 3C).

### Study III: Antidepressant-like effects of ketamine in letrozole pre-treated male and female rats

When analyzing all raw data through three-way ANOVAs, significant sex differences emerged in almost all features evaluated (see Supplemental Table S2: effects of Sex). Overall, and similarly to what was described for Study I, female rats showed higher immobility and lower climbing rates in the forced-swim test (at least ##*p* = 0.001 for both measurements; see Supplemental Table S2 and Fig. 3A and 3B), more distance traveled (cm) in the novelty-suppressed feeding test (##*p* = 0.003, Fig. 3C), and higher sucrose intake (g/kg) in the sucrose preference test (### *p* < 0.001, Fig. 3D) than male adult rats. Interestingly, these basal sex differences persisted in time, at least up to 65 days post-treatment, when female rats still showed higher immobility (##*p* = 0.010) and lower climbing rates than males (#*p* = 0.013, Fig. 5; Supplemental Table S2). Because of these noticeable sex differences, the combined effects of letrozole pre-treatment and ketamine treatment were analyzed for each sex separately through two-ways ANOVAs (Supplemental Table S2; Fig. 4 and 5).

**Fig. 4.**
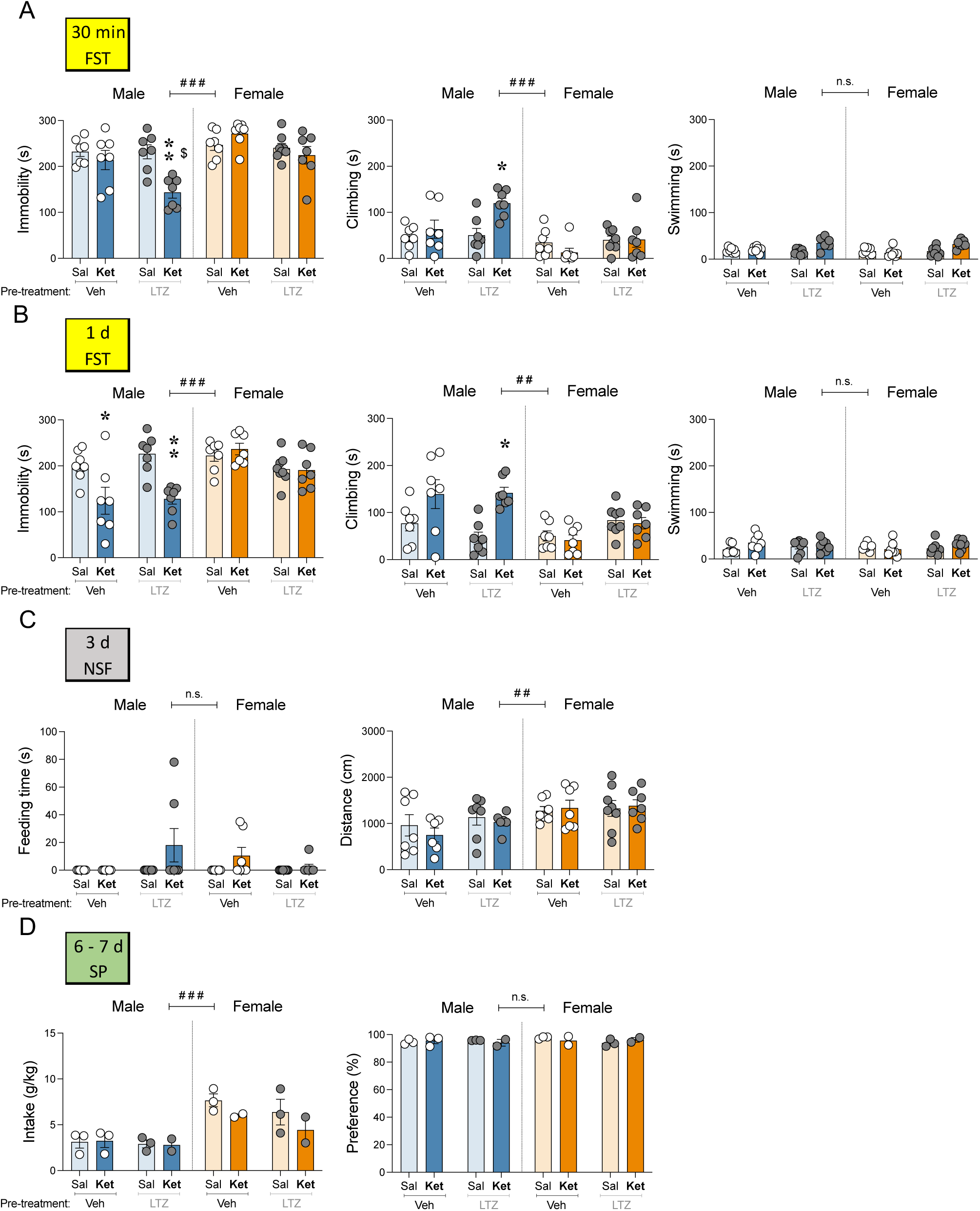
Antidepressant-like effects of ketamine in letrozole pre-treated male and female rats. **A** acute (30 min post-treatment) and **B** repeated (1-day post-treatment) effects exerted by ketamine exposure in male and female rats pre-treated with vehicle or letrozole in the forced-swim test (FST). Data represent mean ± SEM of the time spent (s) immobile, climbing, or swimming. **C** Repeated effects (3-days post-treatment) exerted by ketamine exposure in male and female rats pre-treated with vehicle or letrozole in the novelty-suppressed feeding test (NSF). Data represent mean ± SEM of the feeding time (s), or distance traveled (cm). **D** Repeated effects (6–7 days post-treatment) exerted by ketamine exposure in male and female rats pre-treated with vehicle or letrozole in the sucrose preference test (SP). Data represent mean ± SEM of the sucrose intake (g/kg) or preference (%). **A-D** Individual values are shown for each rat (symbols). Three-way ANOVAs (independent variables: Sex, Pre-Treatment and Treatment) or two-way ANOVAs (independent variables: Pre-Treatment and Treatment) were performed and results are shown in Supplemental Table S2. ^###^*p*<0.001 and ^##^*p*<0.01 when comparing female vs. male rats (effect of Sex). \*\**p*<0.01 and \**p*<0.05 for LTZ-Ket vs. LTZ-Sal and ^$^*p*<0.05 for LTZ-Ket vs. Veh-Ket. Sal: saline; Ket: ketamine; Veh: vehicle; LTZ: letrozole.

**Fig. 5.**
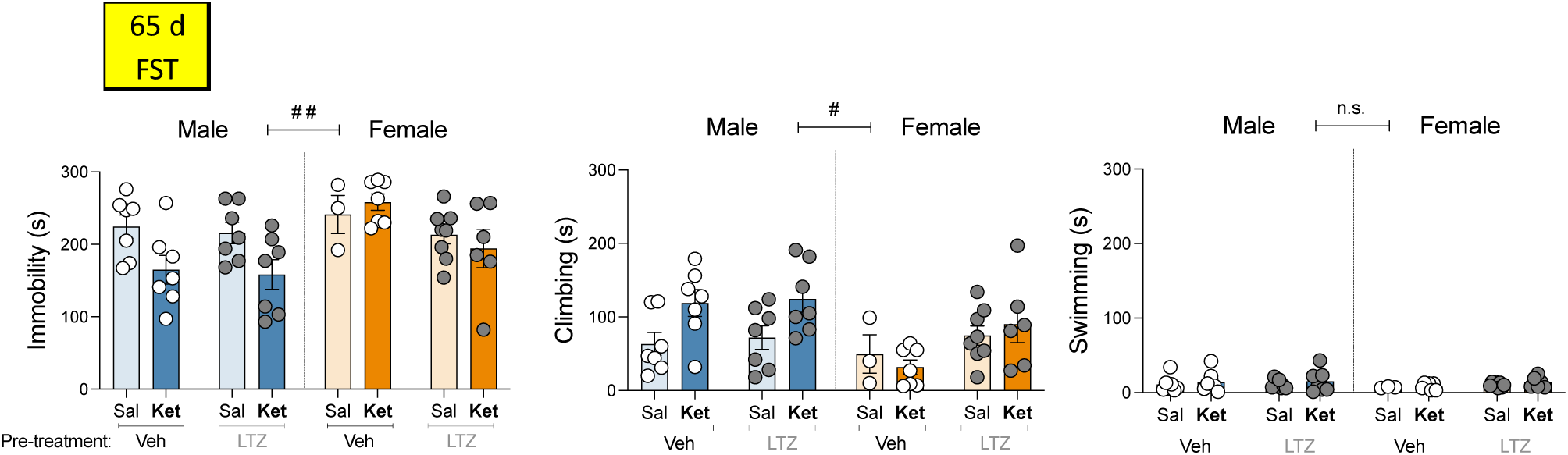
Long-lasting antidepressant-like effects of ketamine or letrozole in rats. Repeated (65-day post-treatment) effects exerted by ketamine exposure in male and female rats pre-treated with vehicle or letrozole in the forced-swim test (FST). Data represent mean ± SEM of the time spent (s) immobile, climbing, or swimming. Three-way ANOVAs (independent variables: Sex, Pre-Treatment and Treatment) or two-way ANOVAs (independent variables: Pre-Treatment and Treatment) were performed and results are shown in Supplemental Table S2. ^###^*p*<0.01 and ^#^*p*<0.05 when comparing female vs. male rats (effect of Sex). Sal: saline; Ket: ketamine; Veh: vehicle; LTZ: letrozole.

The response induced by acute ketamine was measured in the forced-swim test 30 min post a single injection in rats pre-treated or not with letrozole. The results showed that, comparably to what was reported for Study I, acute ketamine did not induce changes in immobility, climbing or swimming behaviors in adult male or female rats (vehicle pre-treated; Fig. 4A). However, in adult male rats pre-treated with letrozole, acute ketamine was capable of inducing an antidepressant-like response, an effect that was not observed in females. Particularly, in male rats pre-treated with letrozole ketamine decreased immobility (-88 ± 22 s, \*\**p* = 0.003 vs. Sal-LTZ; -71 ± 22 s, $*p* = 0.020 vs. Ket-LTZ; Fig. 4A) and increased climbing (+69 ± 20 s, \**p* = 0.013 vs. Sal-LTZ; Fig. 4A). Interestingly, a significant effect of Pre-treatment was observed for female rats (Supplemental Table S2) both for immobility and climbing as measured 1-day post-treatment. This overall antidepressant-like effect induced by letrozole in female rats (i.e., decreased immobility and increased swimming; Fig. 4B) was also observed in Study II following an 8-day treatment paradigm.

Following a repeated paradigm of ketamine, again, and similarly to the results reported in Study I, significant effects of Treatment were present both for immobility and climbing exclusively in male rats as measured 1-day post-treatment (see Supplemental Table S2). Sidak’s *post-hoc* comparisons revealed that ketamine decreased immobility time both in vehicle (-77 ± 27 s, \**p* = 0.047 for Veh-Sal vs. Veh-Ket:) and letrozole (-98 ± 26 s, \*\**p* = 0.007 for LTZ-Sal vs. LTZ-Ket) pre-treated male rats (Fig. 4B). In the novelty-suppressed feeding test (as measured 3 days post-treatment), no significant effects of Treatment were detected (see Supplemental Table S2) when measuring feeding time (s) or distance traveled (cm), independently of Pre-treatment, in male and female rats (Fig 4C and Supplemental Table S2). Similarly, no significant effects of Treatment or Pre-treatment (Supplemental Table S2) were observed for 1% sucrose intake (g/kg) or preference (%) in the sucrose preference test in male and female rats 6–7 days post-treatment (Fig. 4D).

Finally, the potential long-term antidepressant-like effect of these drugs was evaluated in the forced-swim test 65 days post-treatment for each sex separately. Interestingly, the results in male rats showed an overall effect of Treatment, demonstrating a persistent antidepressant-like response of ketamine (vs. saline-male treated rats, and independently of Pre-treatment) by decreasing immobility (-58 ± 25 s; *p* = 0.003) and increasing climbing (+54 ± 24 s; *p* = 0.004) in the forced-swim test (see Supplemental Table S2 and Fig. 5). Moreover, in female rats there was on overall effect of Pre-treatment (see Supplemental Table S2; Fig. 5), suggesting a persistent antidepressant-like effect induced by letrozole (vs. vehicle-female treated rats) both for immobility (-46 ± 27 s; *p* = 0.028) and climbing (+42 ± 25 s; *p* = 0.038) in the forced-swim test (see Supplemental Table S2, and Fig. 5).

## Discussion

This study proved that estrogen production is key in modulating some sex-specific antidepressant-like responses in Sprague-Dawley rats. While ketamine was efficacious after a repeated paradigm exclusively in male rats, its combination with letrozole, an inhibitor of estrogen biosynthesis, induced a faster efficacy as it was observed right after a single dose. However, although ketamine rendered inefficacious in female rats, decreasing estrogen production with letrozole induced on itself an antidepressant-like response, which paralleled significant increases in testosterone levels, and suggesting a beneficial role for letrozole in females. Interestingly, both the antidepressant-like effects induced by ketamine in male rats and letrozole in female rats persisted over time (i.e., up to 65 days post-treatment) suggesting long-term sex-directed antidepressant-like effects for these drugs.

The present results showed clear and interesting sex differences in the pharmacological actions exerted by ketamine. Particularly, while ketamine showed certain efficacy in male rats (following a repeated treatment of 5 mg/kg during 7 days), female rats were unresponsive to the expected beneficial antidepressant-like effects of ketamine. In line with our results prior data showed a beneficial response after a repeated paradigm with low doses of ketamine (5, 10 and 15 mg/kg) in Wistar adult male rats [31]. Moreover, in adolescent Sprague-Dawley rats, ketamine (5 mg/kg) induced an acute antidepressant-like effect (after a single dose) which was sex- and stress-dependent, since it was observed in naïve male rats and in maternally-deprived female rats [9]. Other acute doses tested of ketamine also worked in male adult rats (5, 10, 15 mg/kg) [6, 32]. However, and contrarily to our results, other studies showed that female rats were more sensitive to the antidepressant-like effects of ketamine than males, as they responded to lower doses (e.g., 2.5 mg/kg) in the forced-swim test (see review by [1]). These inconsistent results go along with prior clinical and preclinical studies reporting some sex-related differential responses for ketamine’s actions, with not a clear outcome in terms of which sex benefits the most from this particular treatment (reviewed by [1]), and suggested that the disparities reported could be due to differences in the dose/treatment regimens, in the behavioral tests utilized to score the results, and/or in the animals or particular strains used, which overall introduce a higher degree of variability in the findings.

Sex hormones appear to be critical mediators of the antidepressant-like response of ketamine and needs further research, especially because it is overlooked in much of the available literature. For example, gonadal hormones have been implicated in the possible modulation of the antidepressant-like effect induced by ketamine at low doses, since the response was abolished in ovariectomized female rats and also recovered after receiving a hormone replacement treatment [6, 33]. In this context, the present study aimed at evaluating whether inhibiting the synthesis of estrogens through the use of letrozole, an aromatase inhibitor, would improve the antidepressant-like response of ketamine in male rats and/or would allow for ketamine to induce an antidepressant-like effect in female rats. To do so, we first characterized the response induced by letrozole alone, both at the blood hormonal level and as a potential antidepressant in the forced-swim test, since prior preclinical studies already showed that inhibiting the synthesis of estrogen either with letrozole [19, 34], or formestane [35], could induce an antidepressant like-effect in female rats, and increase hippocampal neurogenesis (i.e., as a neuroplasticity sign of an antidepressant-like action; [36]). Our results showed that a sustained treatment with letrozole (8 days) significantly increased the blood levels of testosterone in female rats as expected, since blocking ovarian aromatization results in a high accumulation of testosterone in females. In male rats, however, the levels did not change, since a high amount of testosterone is expected to be metabolized to non-aromatizable androgens, such as dihydrotestosterone [34]. Interestingly, this same treatment with letrozole, in line with prior results [19, 34], induced an antidepressant-like effect exclusively in female rats, which parallel the increased in testosterone levels observed 1 h post-treatment. Remarkably, the repeated administration of testosterone was previously shown to induce an antidepressant-like response in the forced-swim test [37] and to increase hippocampal neurogenesis (see review by [38]), suggesting a role for this hormone in the beneficial effects induced by letrozole in female rats.

In regards to the antidepressant-like effects of ketamine in letrozole pre-treated male and female rats, the results showed, as hypothesized, that letrozole improved the response of ketamine in male rats (i.e., effect observed right after a single dose in the forced swim test). Moreover, and replicating the data presented before, a repeated paradigm of ketamine was needed to induce an antidepressant-like response in vehicle-pretreated rats, as observed 1-day post-treatment. At this time point, combining letrozole and ketamine did not show a higher response than just ketamine, suggesting probably a maximum effect in the assay evaluated. However, letrozole did not improve the lack of response induced by ketamine in female rats, which still rendered inefficacious, but once again induced an overall antidepressant-like effect. The mechanism by which letrozole may potentiate the antidepressant-like effects of ketamine in adult male rats is unknown. One could speculate that letrozole could either be acting synergistically on the same molecular pathways targeted by ketamine or in additional molecular pathways, producing the enhanced physiological response observed. Moreover, and as discussed in the context of the beneficial effects of letrozole observed in females, it is possible that the increase in testosterone that accompanies aromatase inhibition might be responsible for the improved antidepressant-like response (i.e., [37–38]) induced by ketamine. Moreover, the inhibition of estrogen biosynthesis by letrozole could be affecting the pharmacokinetic and pharmacodynamic profiles of ketamine, since hormonal fluctuations are postulated as the main cause of variability in these processes for any drug [39]. Further studies will be centered in evaluating the molecular mechanisms whereby letrozole increased the efficacy of ketamine in adult male rats.

Finally, one of the most remarkable results presented in this study is the fact that, both the antidepressant-like effects induced by ketamine in adult male rats or letrozole in adult female rats persisted over a long time, since they were still present up to 65 days post-treatment, suggesting long-term sex-directed benefits for these drugs. This is relevant in the context that no prior studies have evaluated persistent effects for so long. For example, a single dose of ketamine induced an acute antidepressant-like effect that long-lasted up to 7 days [40], but nothing further has been evaluated after that time. Moreover, no studies have described the antidepressant-like effects of letrozole across time. Therefore, these findings might be the first ones demonstrating persistent and long-lasting effects for ketamine and letrozole up to 65 days post-treatment, proving a great therapeutical potential in a sex-driven manner (i.e., ketamine for male rats and letrozole for females).

Overall, these findings have proven clear sex-differences in the antidepressant-like response of ketamine in rats, with male rats being more responsive to its beneficial effects proving long-term efficacy. Moreover, inhibiting the biosynthesis of estrogen with letrozole improved the efficacy of ketamine in male rats by advancing the response right after a single dose, in comparison to the repeated paradigm needed in normal conditions. Also, ketamine did not show an antidepressant-like potential in female rats, at least with the conditions tested, however, letrozole induced a remarkable antidepressant-like response in female rats, which persisted in time. Based on this study, ketamine should be utilized adhering to sex-specific considerations. Moreover, inhibiting the biosynthesis of estrogen with letrozole presents itself as a great pharmacological target, since it improved ketamine’s efficacy in male rats and induced on itself a promising response in female rats. The production of estrogen and/or the accumulation of testosterone are key players in modulating some sex-specific antidepressant-like responses in Sprague-Dawley rats.

## Acknowledgments

Research was funded by PID2020-118582RB-I00 (MCIN/AEI/10.13039/501100011033); PDR2020/14 (Comunitat Autònoma de les Illes Balears through the Direcció General de Política Universitària i Recerca with funds from the Tourist Stay Tax Law ITS 2017–006) to MJG-F. The program JUNIOR (IdISBa, GOIB) supported SL-C’s salary.

