## Supplemental Table for "Evaluating the role of inhibiting the biosynthesis of estrogens in the sex-specific antidepressant-like effects of ketamine in rats"

### Supplemental Tables

**Supplemental Table S1. Antidepressant-like effects induced by ketamine or letrozole in adult male and female rats.** This table represents the two-way ANOVAs analyses (F(DFn, DFd), *p* value for data represented in Fig. 2 (Study I) and 3 (Study II). Green-shadow boxes represent statistically significant comparisons.

| **Study I: Sex differences in the antidepressant-like effects of ketamine** | | | |
|---|---|---|---|
| **Fig. 2A. 30 min FST** | **Sex** | **Treatment (Sal vs. Ket)** | **Sex x Treatment** |
| Immobility (s) | F(1,28) = 3.68; *p* = 0.065 | F(1,28) = 0.34; *p* = 0.565 | F(1,28) = 2.03; *p* = 0.165 |
| Climbing (s) | F(1,28) = 5.19; #*p* = 0.031 | F(1,28) = 0.01; *p* = 0.970 | F(1,28) = 1.64; *p* = 0.211 |
| Swimming (s) | F(1,28) = 0.15; *p* = 0.704 | F(1,28) = 4.14; *p* = 0.052 | F(1,28) = 1.53; *p* = 0.226 |
| **Fig. 2B. 1 d FST** | **Sex** | **Treatment (Sal vs. Ket)** | **Sex x Treatment** |
| Immobility (s) | F(1,27) = 0.26; *p* = 0.614 | F(1,27) = 6.34; *p* = 0.018 | F(1,27) = 6.09; *p* = 0.020 |
| Climbing (s) | F(1,27) = 0.22; *p* = 0.642 | F(1,27) = 2.58; *p* = 0.120 | F(1,27) = 5.58; *p* = 0.026 |
| Swimming (s) | F(1,27) = 0.33; *p* = 0.569 | F(1,27) = 5.77; *p* = 0.023 | F(1,27) = 1.57; *p* = 0.221 |
| **Fig. 2C. 3 d NSF** | **Sex** | **Treatment (Sal vs. Ket)** | **Sex x Treatment** |
| Feeding time (s) | F(1,28) = 0.30; *p* = 0.587 | F(1,28) = 0.48; *p* = 0.492 | F(1,28) = 0.03; *p* = 0.856 |
| Distance (cm) | F(1,26) = 6.76; #*p* = 0.015 | F(1,26) = 0.13; *p* = 0.720 | F(1,26) = 0.37; *p* = 0.550 |
| **Fig. 2D. 6-7 d SP** | **Sex** | **Treatment (Sal vs. Ket)** | **Sex x Treatment** |
| Intake (g/kg) | F(1,28) = 46.07; ###*p* < 0.001 | F(1,28) = 0.20; *p* = 0.654 | F(1,28) = 3.40; *p* = 0.076 |
| Preference (%) | F(1,28) = 0.09; *p* = 0.767 | F(1,28) = 1.55; *p* = 0.224 | F(1,28) = 1.33; *p* = 0.259 |
| **Study II: Sex differences in the antidepressant-like effects of letrozole** | | | |
| **Fig.3A. 1 h ELISA** | **Sex** | **Pre-treatment (Veh vs. LTZ)** | **Sex x Pre-treatment** |
| Testosterone (ng/ml) | F(1,20) = 10.81; ##*p* = 0.004 | F(1,20) = 0.04; *p* = 0.835 | F(1,20) = 1.43; *p* = 0.247 |
| **Fig.3B. 1 h FST** | **Sex** | **Pre-treatment (Veh vs. LTZ)** | **Sex x Pre-treatment** |
| Immobility (s) | F(1,28) = 25.36; ###*p* < 0.001 | F(1,28) = 0.06; *p* = 0.801 | F(1,28) = 4.52; *p* = 0.043 |
| Climbing (s) | F(1,28) = 21.28; ###*p* < 0.001 | F(1,28) = 0.09; *p* = 0.763 | F(1,28) = 1.69; *p* = 0.212 |
| Swimming (s) | F(1,28) = 1.32; *p* = 0.261 | F(1,28) = 0.06; *p* = 0.811 | F(1,28) = 8.17; *p* = 0.008 |
| **Fig. 3C. 1 d FST** | **Sex** | **Pre-treatment (Veh vs. LTZ)** | **Sex x Pre-treatment** |
| Immobility (s) | F(1,28) = 10.59; ##*p* = 0.003 | F(1,28) = 0.58; *p* = 0.454 | F(1,28) = 0.17; *p* = 0.682 |
| Climbing (s) | F(1,28) = 12.82; ##*p* = 0.001 | F(1,28) = 0.66; *p* = 0.425 | F(1,28) = 0.09; *p* = 0.766 |
| Swimming (s) | F(1,28) = 0.30; *p* = 0.590 | F(1,28) = 0.01; *p* = 0.979 | F(1,28) = 0.29; *p* = 0.595 |

**Supplemental Table S2. Antidepressant-like effects of ketamine in letrozole pre-treated male and female rats.** This table represents the three or two-way ANOVAs analyses (F(DFn, DFd), *p* value for data represented in Fig. 4 and 5 (Study III). Green-shadow boxes represent statistically significant comparisons.

| **Study III: Antidepressant-like effects of ketamine in letrozole pre-treated male and female rats** | | | |
|---|---|---|---|
| **Fig. 4A. 30 min FST** | **Sex** | **Pre-treatment (Veh vs. LTZ)** | **Treatment (Sal vs. Ket)** |
| Immobility (s) | F(1,49) = 16.23; ###*p* < 0.001 | F(1,49) = 9.57; *p* = 0.003 | F(1,49) = 5.93; *p* = 0.019 |
| Climbing (s) | F(1,49) = 17.06; ###*p* < 0.001 | F(1,49) = 6.52; *p* = 0.014 | F(1,49) = 3.19; *p* = 0.080 |
| Swimming (s) | F(1,49) = 1.20; *p* = 0.279 | F(1,49) = 11.09; *p* = 0.002 | F(1,49) = 12.90; *p* < 0.001 |
| **Fig. 4B. 1 d FST** | **Sex** | **Pre-treatment (Veh vs. LTZ)** | **Treatment (Sal vs. Ket)** |
| Immobility (s) | F(1,49) = 12.96; ###*p* < 0.001 | F(1,49) = 1.02; *p* = 0.318 | F(1,49) = 12.83; *p* < 0.001 |
| Climbing (s) | F(1,49) = 11.52; ##*p* = 0.001 | F(1,49) = 0.83; *p* = 0.366 | F(1,49) = 10.25; p = 0.002 |
| Swimming (s) | F(1,49) = 0.59; *p* = 0.447 | F(1,49) = 0.15; *p* = 0.696 | F(1,49) = 1.71; *p* = 0.197 |
| **Fig. 4C. 3 d NSF** | **Sex** | **Pre-treatment (Veh vs. LTZ)** | **Treatment (Sal vs. Ket)** |
| Feeding time (s) | F(1,49) = 0.16; *p* = 0.688 | F(1,49) = 0.52; *p* = 0.472 | F(1,49) = 5.19; *p* = 0.027 |
| Distance (cm) | F(1,47) = 10.08; ##*p* = 0.003 | F(1,47) = 1.51; *p*= 0. 225 | F(1,47) = 0.19; *p* = 0.665 |
| **Fig. 4D. 6-7 d SP** | **Sex** | **Pre-treatment (Veh vs. LTZ)** | **Treatment (Sal vs. Ket)** |
| Intake (g/kg) | F(1,13) = 23.55; ###*p* < 0.001 | F(1,13) = 1.9; *p* = 0.190 | F(1,13) = 2.0; *p* = 0.182 |
| Preference (%) | F(1,13) = 0.53: *p* = 0.480 | F(1,13) = 0.60; *p* = 0.454 | F(1,13) = 0.03; *p* = 0.869 |
| **Fig. 5A. 65 d FST** | **Sex** | **Pre-treatment (Veh vs. LTZ)** | **Treatment (Sal vs. Ket)** |
| Immobility (s) | F(1,44) = 7.25; ##*p* = 0.010 | F(1,44) = 4.07; *p* = 0.049 | F(1,44) = 4.97; *p* = 0.031 |
| Climbing (s) | F(1,44) = 6.76; #*p* = 0.013 | F(1,44) = 3.70; *p* = 0.061 | F(1,44) = 4.35; *p* = 0.043 |
| Swimming (s) | F(1,44) = 1.11; *p* = 0.301 | F(1,44) = 0.75; *p* = 0.392 | F(1,44) = 1.18; *p* = 0.283 |
| **Study III: Antidepressant-like effects of ketamine in letrozole pre-treated male rats** | | | |
| **Fig. 4A. 30 min FST** | **Pre-treatment (Veh vs. LTZ)** | **Treatment (Sal vs. Ket)** | **Pre-treatment x Treatment** |
| Immobility (s) | F (1,24) = 5.37; *p* = 0.029 | F (1,24) = 12.06; *p* = 0.002 | F(1,24) = 5.24; *p* = 0.031 |
| Climbing (s) | F(1,24) = 4.42; *p* = 0.043 | F(1,24) = 9.16; *p* = 0.006 | F(1,24) = 3.43; *p* = 0.076 |
| **Fig. 4B. 1 d FST** | **Pre-treatment (Veh vs. LTZ)** | **Treatment (Sal vs. Ket)** | **Pre-treatment x Treatment** |
| Immobility (s) | F (1,24) = 0.61; *p* = 0.441 | F (1,24) = 21.71; *p* < 0.001 | F(1,24) = 0.32; *p* = 0.574 |
| Climbing (s) | F (1,24) = 0.56; *p* = 0.463 | F (1,24) = 16.59; *p* < 0.001 | F(1,24) = 0.78; *p* = 0.385 |
| **Fig. 4C. 3 d NSF** | **Pre-treatment (Veh vs. LTZ)** | **Treatment (Sal vs. Ket)** | **Pre-treatment x Treatment** |
| Distance (cm) | F(1,22) = 1.70; *p* = 0.205 | F(1,22) = 0.82; *p* = 0.374 | F(1,22) = 0.09; *p* = 0.769 |
| **Fig. 4D. 6-7 d SP** | **Pre-treatment (Veh vs. LTZ)** | **Treatment (Sal vs. Ket)** | **Pre-treatment x Treatment** |
| Intake (g/kg) | F(1,7) = 0.27; *p* = 0.621 | F(1,7) = 0.00; *p* > 0.999 | F(1,7) = 0.03; *p* = 0.864 |

| **Fig. 5A. 65 d FST** | **Pre-treatment (Veh vs. LTZ)** | **Treatment (Sal vs. Ket)** | **Pre-treatment x Treatment** |
|---|---|---|---|
| Immobility (s) | F (1,24) = 0.19; *p* = 0.671 | F (1,24) = 10.61; *p* = 0.003 | F(1,24) = 0.01; *p* = 0.950 |
| Climbing (s) | F (1,24) = 0.17; *p* = 0.678 | F (1,24) = 10.24; *p* = 0.004 | F(1,24) = 0.01; *p* = 0.927 |
| **Study III: Antidepressant-like effects of ketamine in letrozole pre-treated female rats** | | | |
| **Fig. 4A. 30 min FST** | **Pre-treatment (Veh vs. LTZ)** | **Treatment (Sal vs. Ket)** | **Pre-treatment x Treatment** |
| Immobility (s) | F(1,25) = 4.16; *p* = 0.052 | F(1,25) = 0.11; *p* = 0.743 | F(1,25) = 2.44; *p* = 0.131 |
| Climbing (s) | F(1,25) = 2.10; *p* = 0.159 | F(1,25) = 0.75; *p* = 0.394 | F(1,25) = 0.89; *p* = 0.355 |
| **Fig. 4B. 1 d FST** | **Pre-treatment (Veh vs. LTZ)** | **Treatment (Sal vs. Ket)** | **Pre-treatment x Treatment** |
| Immobility (s) | F(1,25) = 8.37; *p* = 0.008 | F(1,25) = 0.23; *p* = 0.638 | F(1,25) = 0.40; *p* = 0.533 |
| Climbing (s) | F(1,25) = 9.44; *p* = 0.005 | F(1,25) = 0.46; *p* = 0.502 | F(1,25) = 0.02; *p* = 0.903 |
| **Fig. 4C. 3 d NSF** | **Pre-treatment (Veh vs. LTZ)** | **Treatment (Sal vs. Ket)** | **Pre-treatment x Treatment** |
| Distance (cm) | F(1,25) = 0.12; *p* = 0.730 | F(1,25) = 0.16; *p* = 0.692 | F(1,25) = 0.01; *p* = 0.989 |
| **Fig. 4D. 6-7 d SP** | **Pre-treatment (Veh vs. LTZ)** | **Treatment (Sal vs. Ket)** | **Pre-treatment x Treatment** |
| Intake (g/kg) | F(1,6) = 1.56; *p* = 0.258 | F(1,6) = 2.49; *p* = 0.166 | F(1,6) = 0.02; *p* = 0.899 |
| **Fig. 5A. 65 d FST** | **Pre-treatment (Veh vs. LTZ)** | **Treatment (Sal vs. Ket)** | **Pre-treatment x Treatment** |
| Immobility (s) | F(1,20) = 5.62; *p* = 0.028 | F(1,20) = 0.01; *p* = 0.960 | F(1,20) = 0.85; *p* = 0.367 |
| Climbing (s) | F(1,20) = 4.962; *p* = 0.038 | F(1,20) = 0.01; *p* = 0.951 | F(1,20) = 0.78; *p* = 0.387 |
